# Impact of vitamin B12 on rhamnose metabolism, stress defense and in-vitro virulence of *Listeria monocytogenes*

**DOI:** 10.1101/2021.08.26.457850

**Authors:** Zhe Zeng, Lucas M. Wijnands, Sjef Boeren, Eddy J. Smid, Richard A. Notebaart, Tjakko Abee

**Affiliations:** Food Microbiology, Wageningen University and Research, Wageningen, The Netherlands; National Institute of Public Health and the Environment, Bilthoven, The Netherlands; Laboratory of Biochemistry, Wageningen University and Research, Wageningen, The Netherlands

## Abstract

*Listeria monocytogenes* is a facultative anaerobe which can cause a severe food-borne infection known as listeriosis. Rhamnose is a deoxyhexose sugar abundant in a range of environments, including the human intestine, and can be degraded by *L. monocytogenes* in aerobic and anaerobic conditions into lactate, acetate and 1,2-propanediol. Our previous study showed that addition of vitamin B12 stimulates anaerobic growth of *L. monocytogenes* on rhamnose due to the activation of bacterial microcompartment (BMC)-dependent 1,2-propanediol utilization with concomitant production of propionate and propanol. Notably, anaerobic propanediol metabolism has been linked to virulence of enteric pathogens including *Salmonella* spp. and *L. monocytogenes*. In this study we investigate the impact of B12 on aerobic and anerobic growth of *L. monocytogenes* on rhamnose, and observed growth stimulation and *pdu* BMC activation only in anaerobically grown cells with B12 added to the medium. Comparative Caco-2 virulence assays, showed that these *pdu* BMC induced cells have significantly higher translocation efficiency compared to aerobically grown cells (without and with added B12) and non-induced anaerobically grown cells, while adhesion and invasion capacity is similar for all cells. Comparative proteomics analysis showed specific and overlapping responses linked to metabolic shifts, activation of stress defense proteins and virulence factors, with RNA polymerase sigma factor SigL; teichoic acids export ATP-binding protein, TagH; DNA repair and protection proteins RadA and DPS; and glutathione synthase GshAB previously linked to activation of virulence response in *L. monocytogenes*, uniquely upregulated in anaerobically rhamnose grown *pdu* BMC induced cells. Our results shed new light into B12 impact on *L. monocytogenes* competitive fitness and virulence.

## 5.1 Introduction

*Listeria monocytogenes* is the causative agent of the foodborne illness listeriosis, a rare but severe disease with a high mortality rate in the immunocompromised, very young, and elderly populations. Furthermore, listeriosis also causes abortions in pregnant women [1,2]. Acquisition of this infection is mainly caused by consumption of contaminated food (predominantly ready-to-eat food) [1, 2]. *L. monocytogenes* is found ubiquitously in natural environments such as soil, silage, groundwater, sewage and vegetation [2, 3]. The food-borne pathogen can grow at low temperatures and can survive a range of environmental stresses, such as low pH and high salt concentrations [3, 4]. Upon ingestion of contaminated food by the host, and following gastric passage, *L. monocytogenes* can bind to epithelial cells in the intestine [3, 5]. Following entry into epithelial cells mediated by Internalin A (InlA) and Internalin B (InlB), *L. monocytogenes* is internalized into the vacuole [3, 6]. After the internalization, *L. monocytogenes* applies listeriolysin O (LLO) encoded by *hly* [5] and two phospholipases, phospholipase A (PlcA) and phospholipase B (PlcB), for vacuolar rupture and escape, which are crucial steps in *L. monocytogenes* pathogenesis [2, 3, 7]. Notably, alternative invasion and translocation routes involving LLO and Listeria adhesion protein (LAP) have been described [8, 9]. All of these features make the transmission and contamination of *L. monocytogenes* a severe concern for the food industry [4, 10, 11].

Recent studies on anaerobic growth of *L. monocytogenes* provided evidence that it has the capacity to form proteinaceous organelles so-called bacterial microcompartments (BMCs) which enable extension of its metabolic repertoire by supporting the utilization of rhamnose-derived 1,2-propanediol and ethanolamine derived from degradation of phospholipids [12-14]. BMCs are self-assembling organelles that consist of an enzymatic core that is encapsulated by a semi-permeable protein shell [14-16]. The separation of the encapsulated enzymes from the cytosol is thought to protect the cell from toxic metabolic intermediates such as aldehydes, and prevent unwanted side reactions [14-16]. These compartmentalized metabolic pathways may confer a competitive advantage to *L. monocytogenes* over commensal gut microbiota, which typically lack BMC operons and thus are unable to utilize the corresponding substrates in anaerobic conditions [13, 17].

Rhamnose is a deoxyhexose sugar abundant in a range of environments including the human intestine, and can be degraded in anaerobic conditions into 1,2-propanediol which can be further metabolized by a range of bacteria including *L. monocytogenes* that contain so-called propanediol utilization (*pdu*) BMCs [13, 18]. The *L. monocytogenes pdu* cluster is organized together with the ethanolamine utilization (*eut*) cluster and cobalamin synthesis (*cob/cbi*) operons, as a single large locus, referred to as the cobalamin-dependent gene cluster (CDGC) [19, 20]. Previous work suggests that the *pdu* pathway contributes to *L. monocytogenes* establishment in the gastrointestinal tract, while the *eut* pathway may be more important during intracellular replication [19, 21]. We recently provided evidence that anaerobically ethanolamine and propanediol grown eut and *pdu* BMC induced *L. monocytogenes* cells showed significantly higher Caco-2 translocation efficacy in trans-well assays compared to non-induced cells.

We previously reported that addition of vitamin B12 enhances anaerobic utilization of rhamnose via *pdu* BMC in *L. monocytogenes* [13], but the impact of B12 on aerobic and anaerobic growth, metabolism and *in vitro* virulence of *L. monocytogenes* cells grown with rhamnose remains to be determined. We therefore assessed the impact of B12 on rhamnose metabolism in aerobically and anaerobically grown *L. monocytogenes*, aligned with metabolic analysis, proteomics and Caco-2 adhesion, invasion and translocation studies. Possible correlations between these parameters are discussed.

## 5.2. Materials and Methods

### 5.2.1 Strain and Culture Conditions

All experiments in this study were carried out with *L. monocytogenes* EGDe aerobically or anaerobically grown at 30°C in defined medium MWB (Modified Welshimer’s broth) [22]. Overnight grown cells in Luria Broth (LB) were washed three times in PBS before inoculation into MWB. MWB was supplemented with 20mM L-rhamnose as sole carbon source with addition of 20nM B12 (defined as *pdu* BMC induced) or without addition of 20nM B12(defined as *pdu* BMC non-induced). Anaerobic conditions were achieved by Anoxomat Anaerobic Culture System with a gas mixture composed of 10% CO_2_, 5% H_2_, 85% N_2_. OD_600_ measurements in MWB were performed every 12 h for 3 days. Plate counting in MWB to quantity Colony Forming Units (CFUs) was performed every 24 h for 3 days. All the growth measurements were performed with three independent experiments with three technical repeats.

### 5.2.2 Analysis of metabolites for Rhamnose metabolism using High Pressure Liquid Chromatography (HPLC)

Samples were taken from the cultures at 0, 24, 48, and 72 h. After centrifugation, the supernatant was collected for the HPLC measurements of rhamnose while the measurement of acetate, lactate, 1,2-propanediol, 1-propanol and propionate was only performed in 72h. The experiment was performed with three biological replicates. Additionally, the standard curves of all the metabolites were measured in the concentrations 0.1, 1, 5, 10, and 50 mM. HPLC was performed using an Ultimate 3000 HPLC (Dionex) equipped with an RI-101 refractive index detector (Shodex, Kawasaki, Japan), an autosampler and an ion-exclusion Aminex HPX-87H column (7.8 mm × 300 mm) with a guard column (Bio-Rad, Hercules, CA). As the mobile phase 5 mM H_2_SO_4_ was used at a flow rate of 0.6 ml/min, the column was kept at 40°C. The total run time was 30 min and the injection volume was 10 μl. All the HPLC measurements were performed with three independent experiments with three technical repeats as described before [13].

### 5.2.3 Proteomics

*L. monocytogenes* EGDe cultures were aerobically or anaerobically grown at 30°C in MWB with 20 mM rhamnose adding 20nM B12 or not. Samples were collected at 48 h of the inoculation and processed as described before [13]. The mass spectrometry proteomics data have been deposited to the ProteomeXchange Consortium via the PRIDE [23] partner repository with the dataset identifier PXD025734 for *pdu* BMC induced and *pdu* BMC non-induced conditions.

### 5.2.4 Caco-2 adhesion, invasion and translocation

Cultures of Caco-2 cells (human intestinal epithelial cells, ATCC HTB-37), production of differentiated cells in 12-well plates were carried out as described before [24, 25]. *L. monocytogenes* overnight grown cells used for adhesion, invasion and translocation were normalized to get the concentration of 8.4 ± 0.1 log CFU/ml.

For adhesion and invasion experiments, Caco-2 with the inoculum 1.6 × 105 cells/well were seeded into the 12-well tissue culture plates (Corning Inc. ID 3513). Inoculated 12-well plates and were incubated for 12–14 days with the medium refreshing every 2 days at 37 °C to establish a confluent monolayer of cells. Adhesion and invasion experiments were started by inoculation with 40 μl of late exponential phase cells of Rhamnose *pdu* BMC induced and non-induced *L. monocytogenes* EGDe resulting in an final inoculum of approximate 7.41 log CFU/well. Then the 12 well plates were centrifuged for 1 min at 175 ×g to create a proximity between the Caco-2 and *L. monocytogenes* cells.

For adhesion and invasion enumeration, after 1h anaerobic incubation without gentamicin, *L. monocytogenes* cells that have not adhered to Caco-2 cells were removed by washing three times with PBS buffer. Half of the wells containing Caco-2 cells were lysed with 1 ml of 1% v/v Triton X-100 in PBS and serially diluted in PBS for quantification of the number of adhered and invaded *L. monocytogenes* cells. The other half of the wells of the Caco-2 cells was subsequently incubated anaerobically for 3 h with 0.3% gentamicin (50 µg/ml, Gibco) to eliminate all extracellular *L. monocytogenes* cells. Thereafter, gentamycin containing medium was removed by washing three times with PBS buffer. The Caco-2 cells were lysed with 1 ml of 1% v/v Triton X-100 in PBS and serially diluted in PBS for quantification of the number of invaded *L. monocytogenes* EGDe cells.

For translocation experiments, 0.8 × 105 Caco-2 cells per mL were seeded ThinCert PET inserts Griener Bio-One 665640) for 12–14 day differentiation. Prior to the translocation assay wells and inserts were washed three times with PBS, and placed in TCM without gentamycin and fetal bovine serum. Translocation was started by adding 20 µl of late exponential cells of Rhamnose *pdu* BMC induced and non-induced *L. monocytogenes* EGDe into the inserts, resulting in an inoculum of approximately 7.17 log CFU/well. After centrifuging the Transwell plates for 1 min at 175x g, the plates were incubated 2 hrs anaerobically. After incubation, the inserts were removed with a sterile forceps and discarded. The contents in the wells was collected for quantification of the translocated number of *L. monocytogenes* EGDe cells.

### 5.2.5 Venn diagram analysis and STRING networks analysis

The protein IDs of significantly changed proteins from Supplementary Table 1, 2 and 3 were uploaded to the BioVenn online server [26] taking the default setting to generate Venn diagrams. The analysis of Venn diagram and details of overlap upregulated proteins are shown in Supplementary Table 4. Overlapping proteins from the Venn diagram were transferred to the STRING online server[27] for multiple proteins analysis of functional interaction using sources such as co-expression, genomic neighbourhood and gene fusion.

Statistical analyses were performed in Prism 8.0.1 for Windows (GraphPad Software). As indicated in the figure legend, Statistical significances are shown in ***, P<0.001; *, P<0.05; ns, P>0.05 with Holm-Sidak T-test.

## 5.3. Results

### 5.3.1 Impact of vitamin B12 on *L. monocytogenes* grown on rhamnose

We first examined the impact of B12 on aerobic or anaerobic growth of *L. monocytogenes* EGDe in MWB defined medium without and with added vitamin B12 (Figure 1). In Figure 1A, *L. monocytogenes* showed a similar aerobic growth in MWB defined medium plus 20mM rhamnose with or without 20mM B12 where OD_600_ reaches a maximum of approximate 0.56 after 72 h. But for anaerobic conditions, in MWB defined medium supplied with 20mM rhamnose OD_600_ reaches a maximum of about 0.37 after 48 h, while in MWB supplied with 20mM rhamnose and 20nM B12 OD_600_ continues to increase after 48 h, reaching a significant higher OD_600_ of 0.51 at 72 h as described before [13]. The growth phenotypes indicate that addition of 20nM of B12 into MWB defined medium plus 20mM rhamnose stimulates anaerobic growth of *L. monocytogenes*, but has no significant effect on aerobic growth which is already enhanced compared to that in anaerobic conditions. Next, utilization of rhamnose in aerobic and anaerobic conditions with or without B12 was quantified. The aerobic growth of *L. monocytogenes* with or without B12 appears similar including the rhamnose degradation capacity, that showed complete consumption of 20mM rhamnose within 72 h (Figure 1B). Complete consumption of rhamnose was also observed in anaerobically grown cells in MWB plus 20mM rhamnose and B12, whereas 3.5 mM of rhamnose was still present at 72 h in MWB plus 20mM rhamnose without added B12. These results indicate that addition of B12 stimulates anaerobic growth and rhamnose metabolism of *L. monocytogenes* EGDe, while no significant impact of B12 was detected with aerobically grown cells.

**Figure 1.**
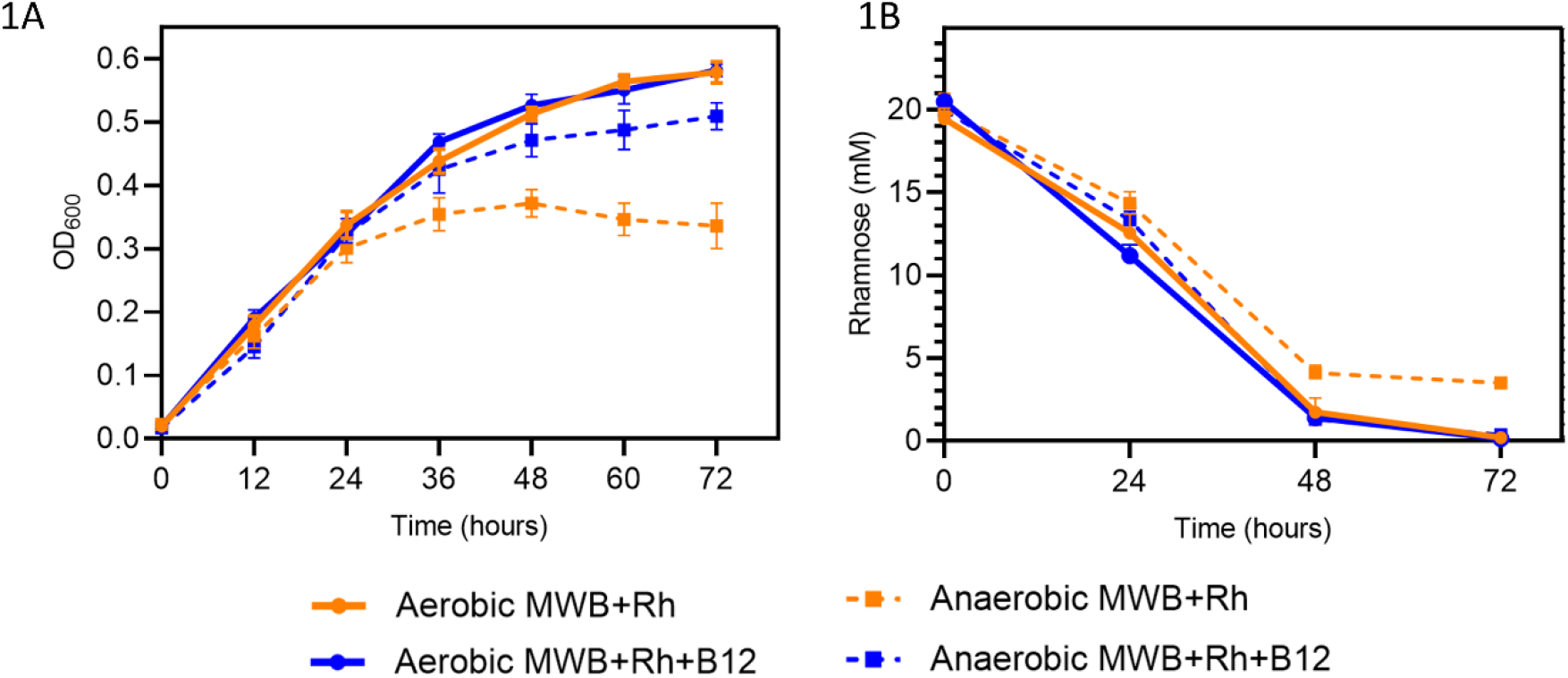
Growth and Rhamnose catabolism of *L. monocytogenes* EGDe in MWB medium with 20mM Rhamnose. **(A)** Impact of vitamin B12 on aerobic or anaerobic growth of *L. monocytogenes* EGDe in MWB medium with 20mM Rhamnose; **(1B)** Rhamnose utilization. Orange lines with closed circles represent *L. monocytogenes* grown in MWB medium with 20 mM Rhamnose, while blue lines with closed circles represent *L. monocytogenes* grown in MWB medium with 20 mM Rhamnose and 20nM B12. Solid lines represent aerobic condition, and striped lines represent anaerobic condition; data published in [13]. Results from three independent experiments are expressed and visualized as means and standard errors.

### 5.3.2 Impact of B12 on aerobic or anaerobic metabolism of rhamnose

Metabolite production in the different growth conditions was analysed by HPLC. As shown in Figure 2, at 72 h, rhamnose was degraded by *L. monocytogenes* EGDe into acetate, lactate and 1,2-propanediol in anaerobic and aerobic conditions with or without B12, albeit to different levels. Metabolites following growth in MWB medium plus 20mM rhamnose with and without B12 in aerobic conditions, showed no significant differences in acetate, lactate and 1,2-propanediol production, in line with the corresponding similar growth phenotypes and rhamnose utilization in Figure 1. Notably, the type and amount of end products formed during rhamnose utilization in anaerobic conditions with B12 added, are significantly different from all other tested conditions, and include 4.1 mM acetate, 2.3 mM lactate, 1.4 mM 1,2-propanediol, and 3.2 mM propionate and 3.6 mM 1-propanol, indicative of activated *pdu* BMC. These data provide evidence for specific activation of *pdu* BMC in *L. monocytogenes* during growth on rhamnose in anaerobic conditions with B12 added to the medium.

**Figure 2.**
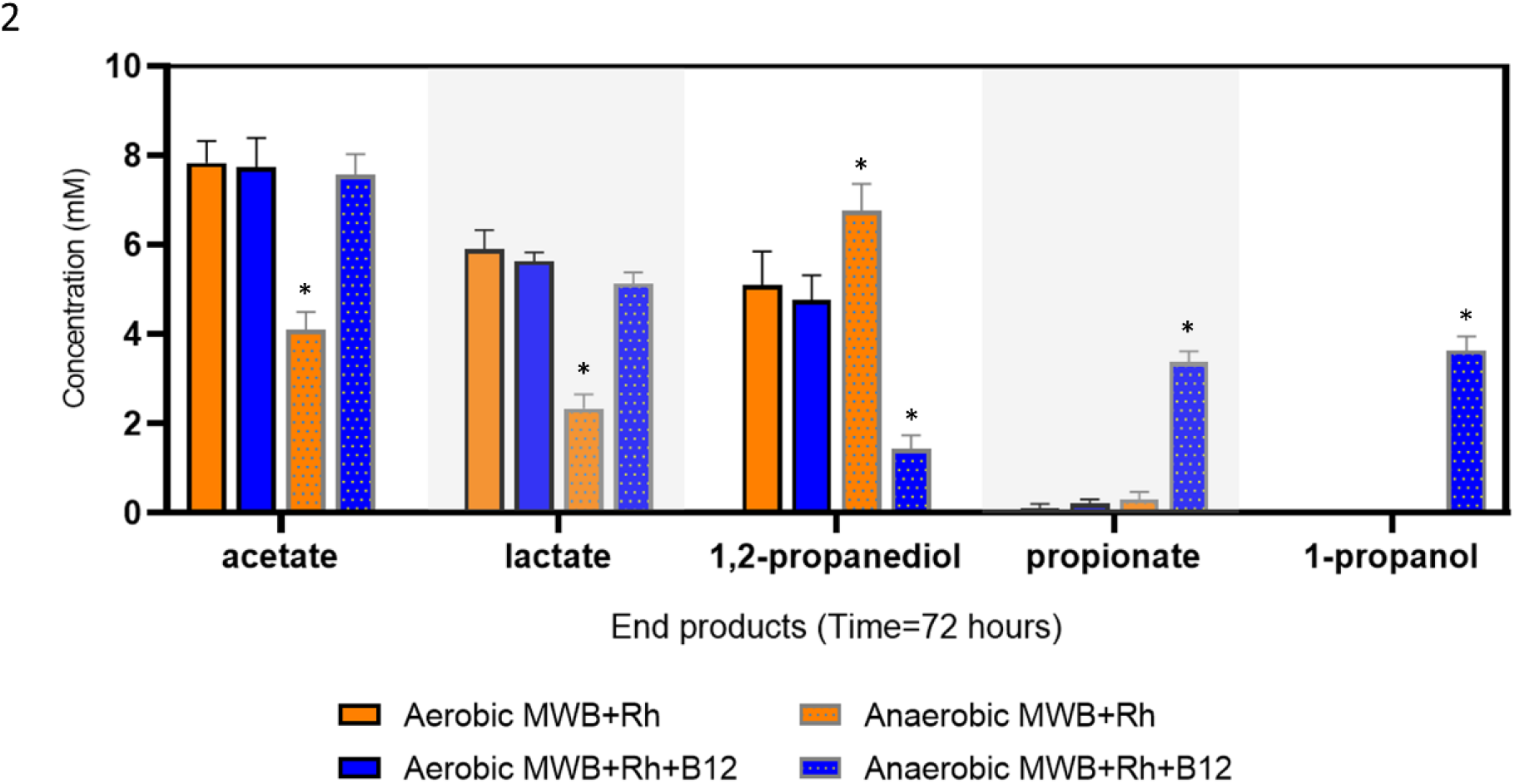
Metabolites from rhamnose metabolism of *L. monocytogenes* EGDe aerobically or anaerobically grown in MWB with 20mM Rhamnose with/without 20nM vitamin B12. Orange bars represent *L. monocytogenes* grown in MWB medium with 20 mM Rhamnose, while blue bars represent *L. monocytogenes* grown in MWB medium with 20 mM Rhamnose and 20nM B12. Metabolites are grouped as indicated in X-axis, successively, acetate, lactate, 1,2-propanediol, propionate and 1-propanol. Bars with black outline represent aerobic condition, and Bars with grey outline and filled pattern represent anaerobic condition; data published in [13]. Results from three independent experiments are expressed and visualized as means and standard errors.

### 5.3.3 Proteomic analysis on the impact of B12 on aerobic or anaerobic growth of *L*.*monocytogenes* on rhamnose

Comparative proteomics analysis identified differentially expressed proteins in cells grown in MWB medium plus 20mM rhamnose with or without B12 in aerobic and anaerobic conditions (Figure 3). Figure 3A shows data for MWB plus 20mM rhamnose in anaerobic condition compared to MWB plus 20 mM rhamnose in aerobic condition. Obviously, 21 proteins out of 23 Pdu proteins (yellow dots encircled in black lines in Figure 3A) in the BMC-dependent 1,2-propanediol utilization cluster, are upregulated in anaerobically grown cells on rhamnose compared to aerobically grown cells (Details in Supplementary Table 1).

**Figure 3.**
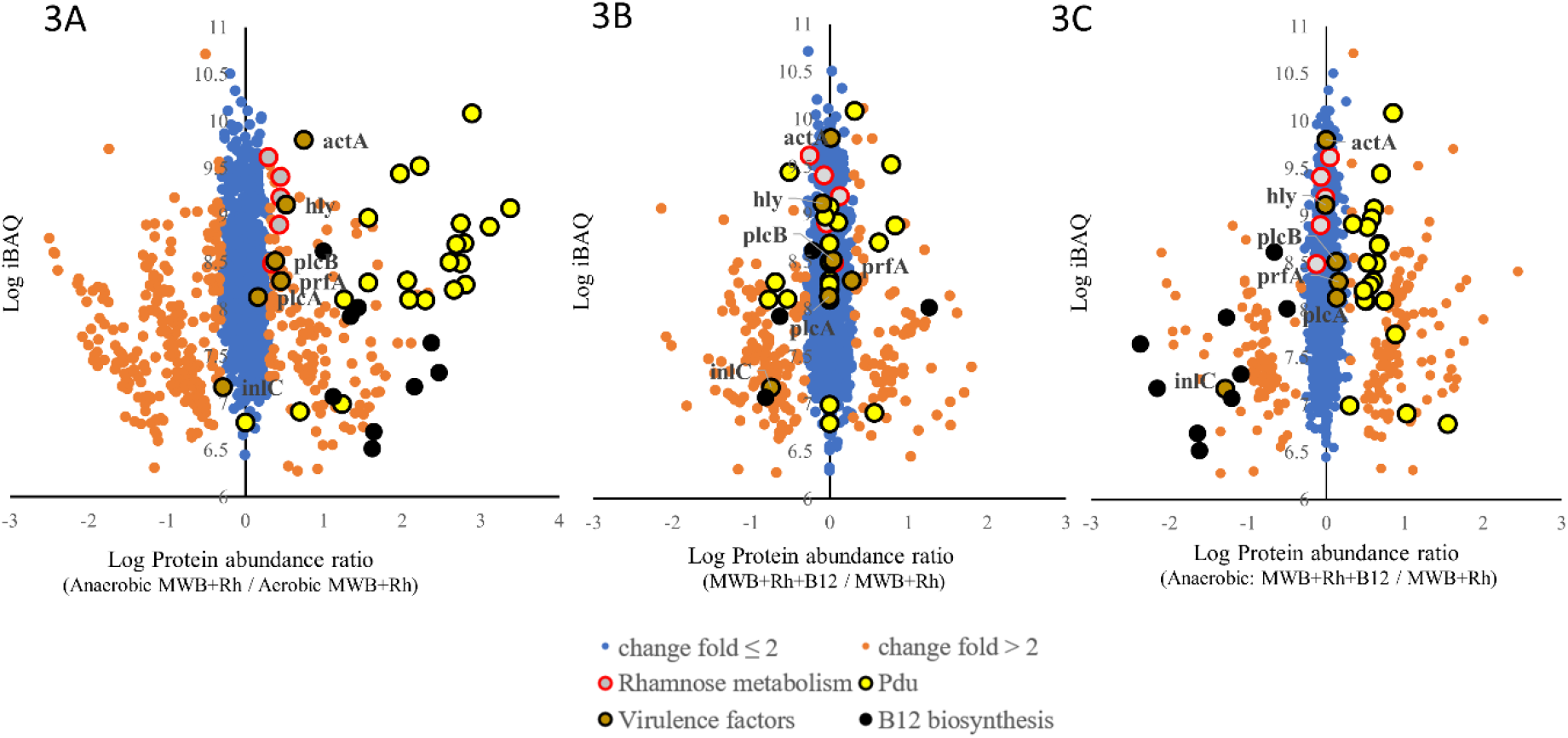
Proteomic analysis of *L. monocytogenes* EGDe aerobically or anaerobically grown in MWB with 20mM Rhamnose with/without 20nM vitamin B12. **(A)** Proteomic ratio plot of *L. monocytogenes* anaerobically grown in MWB plus 20 mM rhamnose compared to *L. monocytogenes* aerobically grown in MWB plus 20 mM rhamnose, (full list in Supplementary Table 1). **(B)** Proteomic ratio plot of *L. monocytogenes* aerobically grown in MWB plus 20 mM rhamnose and 20nM B12 compared to *L. monocytogenes* aerobically grown in MWB plus 20 mM rhamnose, (full list in Supplementary Table 2). **(C)** Proteomic ratio plot of *L. monocytogenes* anaerobically grown in MWB plus 20 mM rhamnose and 20nM B12 compared to *L. monocytogenes* anaerobically grown in MWB plus 20 mM rhamnose, (full list in Supplementary Table 3). Fold change ≤ 2 in blue, fold change > 2 in light orange, proteins in the *pdu* cluster are in yellow, and proteins in the rhamnose cluster are in gray. Virulence factors in Supplementary Table 4 are dark orange. Genes of *cob/cbi* operon for B12 biosynthesis are in black.

Notably, despite expression of Pdu proteins, no production of *pdu* BMC signature metabolites propionate and 1-propanol was detected as shown above (figure 2), in line with our previously reported data [13], that showed only presence of BMCs and activation of *pdu* BMC in anaerobically rhamnose grown cells with added B12. Our data extend previous RNAseq studies that showed slight increase in *pdu* expression in the presence of propanediol, and highest expression when B12 was also present [28, 29]. The lack of *pdu* BMC activation in anaerobic MWB plus rhamnose conditions, suggests that the observed activation of B12 synthesis enzymes does not result in the production of sufficiently high levels of (de novo) B12 to trigger full induction (Figure 3A and Supplementary Table 5).

Comparative analysis of anaerobically versus aerobically grown cells, points to repression of propanediol-induced *pdu* expression in the latter condition. Notably, highlighted virulence factors in the volcano plot (Figure 3 A), show that actin assembly-inducing protein ActA, Listeriolysin O (LLO) encoded by *hly*, Listeriolysin regulatory protein PrfA, and phospholipase C encoded by *clpB*, are slightly upregulated in anaerobic growth compared to aerobic growth of *L. monocytogenes* on rhamnose, while PlcA, phospholipase A, shows no significant difference, and internalin C was repressed. Rha proteins for rhamnose utilization including rhaA, rhaB, rhaD, rhaM and lmo2850 contained in the *rha* cluster, are all slightly upregulated in anaerobic growth compared to aerobic growth of *L. monocytogenes* on rhamnose.

Anaerobic growth of *L. monocytogenes* on rhamnose with added B12 results in increased expression of Pdu proteins (Figure 3C) and presence of BMCs was evidenced by TEM (Shown in our recent study [13]). Rha proteins do not show significant differences, while PrfA, PlcB and ActA are induced, but PlcA, hly and InlC are repressed in MWB plus 20mM rhamnose with B12 compared cells grown without B12 (Details in Supplementary Table 3). Notably, addition of B12 and induction of *pdu* BMC in anaerobic conditions, results in a significant repression of B12 biosynthesis proteins (8 identified B12 proteins, Figure 3C). Reduced expression is also seen in aerobically grown cells, where only 4 B12 biosynthesis proteins are identified (Figure 3B). Combining these results suggests that addition of B12 has a repressive effect on B12 biosynthesis proteins in anaerobically rhamnose grown *L. monocytogenes* EGDe cells, with the B12 biosynthesis proteins largely repressed in aerobically grown cells (Details in Supplementary Table 5).

### 5.3.4 Stress and virulence proteins triggered by B12-activated *pdu* BMC

The Venn diagram provides an overview of differentially expressed proteins in conditions without and with added B12 in aerobically and anaerobically rhamnose grown *L. monocytogenes* (Figure 4). As shown in Figure 4A, 145 proteins (Group X, red pie chart) are upregulated more than two fold in anaerobic MWB plus rhamnose with B12 compared to anaerobic condition without B12, and 81 proteins (Group Y, yellow pie chart) are upregulated more than two fold in aerobic condition with B12 compared to aerobic condition without B12, while 162 proteins (Group Z, blue pie chart) are upregulated more than two fold in anaerobic condition without B12 compared to aerobic condition without B12 (details in Supplementary Table 4). The overlap of group X and group Y contains 25 proteins that are upregulated in in anaerobic and aerobic rhamnose conditions with added B12, with eight of the proteins present also in group Z. The proportion in red group X without overlap with other groups represents 103 proteins upregulated in *pdu* BMC induced cells grown in rhamnose with added B12 (Figure 4B, details in Supplementary Table 4).

**Figure 4.**
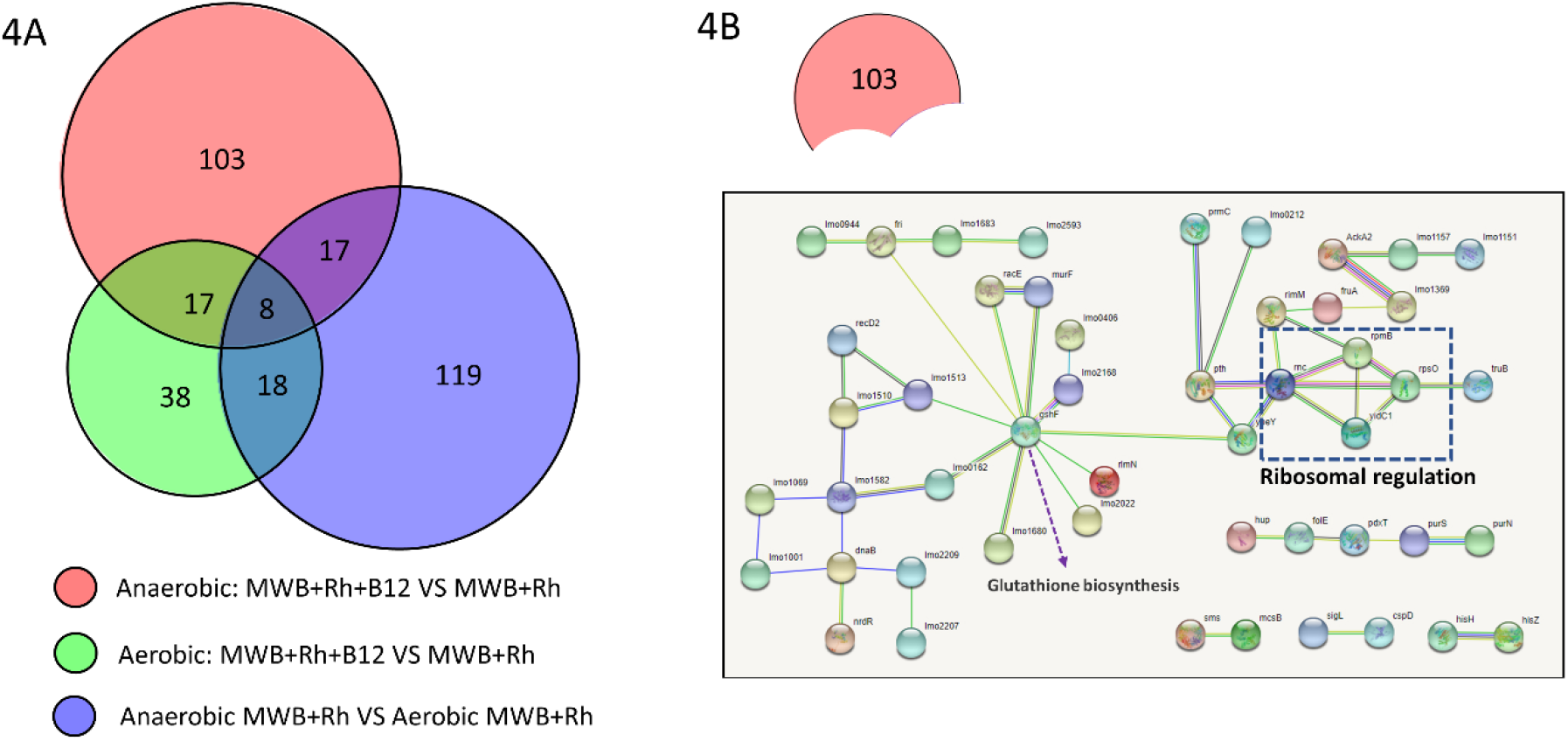
Proteomics analysis of vitamin B12 impact on *L. monocytogenes* EGDe general stress response and virulence factors. **(A)** Venn diagram of overlapping upregulated proteins among three groups. Group X in red: anaerobic growth of *L. monocytogenes* on MWB with 20mM rhamnose and 20nM B12, compared to MWB with 20mM rhamnose without B12; Group Y in yellow: aerobic growth of *L. monocytogenes* on MWB with 20mM rhamnose and 20nM B12 compared MWB with 20mM rhamnose without B12; Group Z in blue: anaerobic growth of *L. monocytogenes* on MWB with 20mM rhamnose without B12 compared aerobic MWB with 20mM rhamnose without B12; **(B)** STRING analysis of proteins specifically upregulated by the addition of B12 in anaerobically grown *L. monocytogenes* in MWB with 20mM rhamnose; Nodes represent proteins, and lines represent interactions, disconnected proteins are not shown in the network (Details in Supplementary Table 4).

STRING protein-protein interaction analysis of these 103 proteins uniquely upregulated in anaerobically rhamnose grown *pdu* BMC induced *L. monocytogenes* cells, points to differential expression of a range of ribosomal and ribosome-associated proteins, including RplB, 50S ribosomal protein L28, RpsO, 30S ribosomal protein S15, Rnc, ribonuclease 3, involved in the processing of primary rRNA, and YbeY, an endoribonuclease involved in late-stage 70S ribosome maturation, while other proteins point to metabolic shifts, activation of stress defense, and links to virulence, such as RNA polymerase sigma factor SigL, Teichoic acids export ATP-binding protein, TagH, DNA repair and protection proteins RadA and DPS, and glutathione synthase GshAB, previously linked to activation of virulence response in *L. monocytogenes* [30-32]. In addition, *pdu* BMC induced cells share a range of other virulence factors also expressed in anaerobic non-induced cells (Supplementary Table 3), including endopeptidase p60 (Iap), (higher expressed in *pdu* BMC cells), Listeriolysin regulatory protein (PrfA), 1-phosphatidylinositol phosphodiesterase (PlcA), Phospholipase C (PlcB), Actin assembly-inducing protein (ActA), Internalin C (InlC) and Internalin B (InlB) (both higher expressed in in non-induced cells), and bi-functional Listeria associated protein (LAP, lmo1634) [2, 3, 8, 33, 34]. Next, *in vitro* virulence assays were performed to compare performance of anaerobic *pdu* BMC induced cells to non-induced anaerobic and aerobic cells.

### 5.3.5 Impact of B12 on Caco-2 cell adhesion, invasion and translocation of *L. monocytogenes* grown aerobically or anaerobically on rhamnose

To address the impact of B12 on *L. monocytogenes in vitro* virulence, we conducted adhesion, invasion and translocation assays with Caco-2 cells using rhamnose *pdu* BMC induced and non-induced cells grown anaerobically and aerobically. As shown in Figure 5, *L. monocytogenes* cells grown anaerobically or aerobically in MWB plus 20mM rhamnose with or without B12 show similar adhesion and invasion efficacy of ∼5.2 log and ∼4.1 log CFU/well, respectively. Strikingly, the ability to translocate Caco-2 cells monolayers in a trans-well system, is significantly higher for *L. monocytogenes* cells grown anaerobically in MWB plus rhamnose with B12 compared to the other cells (∼1 log CFU/well improvement), grown anaerobically without B12 and aerobically without and with B12 (Figure 5). These results indicate similar adhesion and invasion capacity of all four tested cell types, while anaerobically rhamnose grown *pdu* BMC induced *L. monocytogenes* cells show significantly higher translocation efficacy compared to the three non-induced cell types.

**Figure 5.**
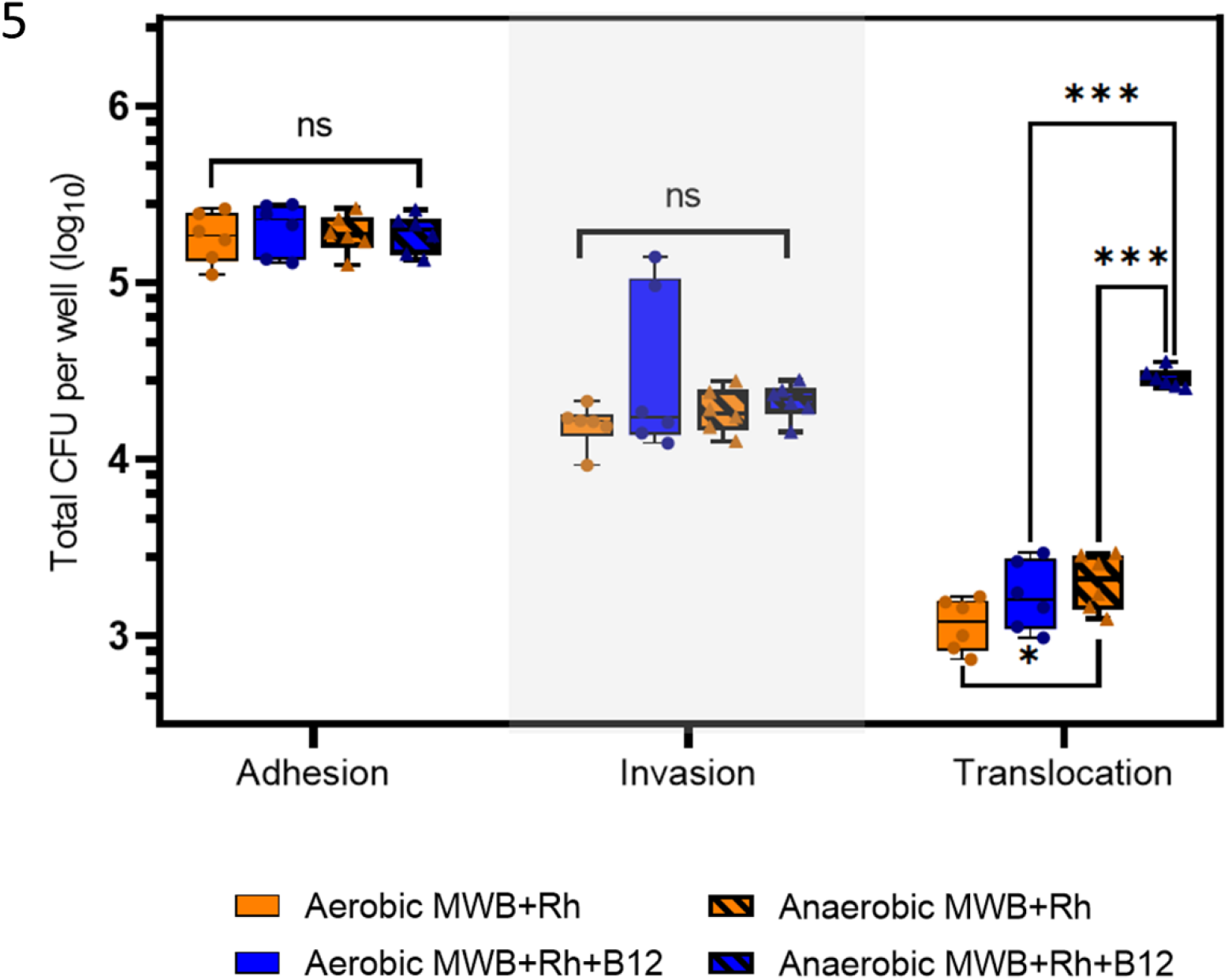
Caco-2 cell assays with *L. monocytogenes* EGDe aerobically or anaerobically grown in MWB with 20mM Rhamnose with/without 20nM vitamin B12. Orange bars represent *L. monocytogenes* grown in MWB medium with 20 mM Rhamnose, while blue bars represent *L. monocytogenes* grown in MWB medium with 20 mM Rhamnose and 20nM B12. Bars without filled pattern represent aerobic conditions, and Bars with filled pattern represent anaerobic conditions. X axis shows three groups of columns with results from adhesion, invasion and translocation assays. Y axis is the total CFU per well in log10 after the assays while initial inputs for adhesion and invasion are 7.41 ± 0.02 in log10 CFU per well and the initial inputs for translocation are 7.17 ± 0.03 in log10 CFU per well. Statistical significance is indicated (*** means P≤0.001; * means P≤0.05; ns means > 0.05 by Holm-Sidak T-test)

## 5.4 Discussion

The results in this study show a unique stimulating effect of B12 supplementation on anaerobic growth of *L. monocytogenes* with rhamnose, linked to *pdu* BMC activation and production of signature metabolites propionate and propanol, while no effect is observed in aerobic conditions. Growth in the latter condition with and without added B12 results in higher biomass production compared to anaerobically grown cells with and without added B12, conceivably due to increased energy generation via respiration [33]. Observed growth stimulation in rhamnose *pdu* BMC induced cells compared to non-induced cells is in line with previously described results [13]. *L. monocytogenes* has to cope with anaerobic conditions encountered during transmission along the food chain, for example in modified atmosphere and vacuum-packed foods, and in the GI tract. Notably, a significant upregulation of Pdu proteins involved in 1,2-propanediol utilization is observed in anaerobic MWB plus 20mM rhamnose grown cells compared to aerobic MWB plus 20mM rhamnose grown cells. This could point to activation of 1,2-propanediol utilization, in line with concomitant increase in expression of B12 synthesis enzymes in these anaerobically rhamnose grown *L. monocytogenes* cells, but HPLC analysis indicates that *pdu* BMC signature metabolites, propionate and 1-propanol are not produced, in line with absence of BMCs in TEM analysis [13]. Addition of B12 to anaerobically rhamnose grown cells results in further, significantly increased, expression of Pdu proteins, and activation of *pdu* BMC as evidenced by the production of propionate and 1-propanol (Figure 2), and presence of BMCs detected by TEM [11,19]. It is conceivable, that B12 dependent activation of *pdu* BMC in anaerobic conditions is not only crucial due to the role of B12 as a cofactor for the signature enzyme PduCDE, a B12-dependent diol dehydratase [20], but also because the PduCDE-B12 complex plays a role in triggering the construction of BMC with the shell proteins thereby encasing *pdu* enzymes, with the respective signature peptide sequences directing enzyme complexes to the correct locations inside the microcompartment [14, 35-37]. Our results with aerobically and anaerobically rhamnose grown cells, indicate that RNAseq data of *pdu* operon, including identification of factors involved in control of gene expression, such as regulator PocR, B12-dependent riboswitch(es) and sRNAs [28, 38], require confirmation by proteomics, TEM and metabolic profiling to enable conclusions about activation of functional *pdu* BMC in *L. monocytogenes*.

B12-dependent activation of *pdu* BMC in *L. monocytogenes* also results in additional cellular responses. Comparative proteome analysis identified 103 proteins uniquely upregulated in *pdu* BMC induced cells with functions in protein synthesis, ribosomal proteins and ribosome-associated proteins, and in metabolism and stress defense, including RNA polymerase sigma factor SigL, Teichoic acids export ATP-binding protein, TagH, DNA repair and protection proteins RadA and DPS, and glutathione (GSH) synthase GshF, a multidomain protein encoded by *gshAB* (Fig 3 and 4, Supplementary Table 2 and 3). GshF mediated production of glutathione was recently linked to activation of virulence response in *L. monocytogenes*, with glutathione acting as an allosteric activator of Listeriolysin regulatory protein PrfA [30]. Notably, a range of virulence factors is expressed both in anaerobic *pdu* BMC induced and non-induced cells (Table 3), including regulatory protein PrfA, phosphatidylinositol phosphodiesterase (PlcA), Phospholipase C (PlcB), listeriolysin O (LLO), Actin assembly-inducing protein (ActA), Internalin C (InlC) and Internalin B (InlB), and bi-functional Listeria associated protein (LAP, lmo1634) [8, 9]. Extracellular cell wall associated LAP, has been suggested to play a role in *L. monocytogenes* interaction with the host by affecting cellular redistribution of epithelial junction proteins enhancing bacterial translocation [8, 9]. Analysis of proteomes from aerobically rhamnose grown cells without and with added B12) (Supplementary Table 1 and 3) also indicate (low level) presence of the virulence factors described above.

Our comparative in vitro virulence assays indicate similar Caco-2 adhesion and invasion capacity of aerobically and anaerobically rhamnose grown cells without and with added B12, while only anaerobically rhamnose grown *pdu* BMC induced *L. monocytogenes* cells show significantly higher Caco-2 translocation efficacy compared to the three other non-induced cell types. These results suggest that anaerobically rhamnose grown *pdu* BMC primed *L. monocytogenes* cells show enhanced transcellular and/or paracellular translocation in the Caco-2 cells trans-well assay. Transcellular translocation involves *L. monocytogenes* binding and internalization into epithelial cells, and proceeds by intracellular replication and/or movement into neighboring epithelial cells by hijacking host cellular machinery via PrfA-activated virulence factors [34, 39, 40]. Based on similar performance in our *L. monocytogenes* EGDe adhesion and invasion studies, with 1 h and 3 h incubation periods, respectively, it is unlikely that enhanced translocation capacity, with a 2 h incubation period, is linked to enhanced transcellular performance of rhamnose *pdu* BMC primed cells. An alternative explanation could be offered by enhanced paracellular translocation of *L. monocytogenes* cells, i.e., passing through an intercellular space between the Caco-2 cells, such as tight junctions [41, 42]. Tight junctions play a major role in maintaining the integrity and impermeability of epithelial cells, including the intestinal barrier [41-43]. Different strategies are used by pathogens aimed at destabilizing tight junctions, and roles for internalin A, LAP and listeriolysin O (LLO) in paracellular translocation of *L. monocytogenes* have been reported [8, 42, 43]. Combined with adapted physiology and metabolic shifts in *pdu* BMC induced cells including activated stress defense and GSH production, action of LAP and/or LLO, may have supported enhanced translocation in the trans-well assay. Notably, our previous results with anaerobically grown *L. monocytogenes* EGDe in LB medium with propanediol or ethanolamine, with and without added B12, showed similar adhesion and invasion capacity, but significantly higher translocation efficacy of respective *pdu* BMC and *eut* BMC induced cells.

We previously provided evidence for *L. monocytogenes* growth stimulation via B12 dependent activation of *pdu* BMC in anaerobically propanediol and rhamnose grown cells [13, 20]. The *pdu* BMC may thus enhance competitive fitness in the intestine, in line with observations that showed reduced *L. monocytogenes* persistence in stool and ileal colonization of female BALB/c mice of a pduD (subunit of propanediol dehydratase) deletion mutant compared to the wild type [44]. Out results obtained in Caco-2 virulence assays may point to an additional impact of (rhamnose) *pdu* BMC activation on *L. monocytogenes* interaction with host intestinal epithelial barrier, more specifically, interaction with tight junctions. Additional experiments are required to confirm our hypothesis and to elucidate underlying mechanisms.

## 5.5 Supplementary Materials

**Supplementary Table 1**. Protein profiling of *L. monocytogenes* EGDe anaerobically grown on MWB with 20mM rhamnose compared to *L. monocytogenes* EGDe aerobically grown on MWB with 20mM rhamnose.

**Supplementary Table 2**. Protein profiling of *L. monocytogenes* EGDe anaerobically grown on MWB with 20mM rhamnose and 20nM B12 compared to MWB with 20mM rhamnose without B12.

**Supplementary Table 3**. Protein profiling of *L. monocytogenes* EGDe aerobically grown on MWB with 20mM rhamnose and 20nM B12 compared to MWB plus 20mM rhamnose without B12

**Supplementary Table 4**. Details of overlap in upregulated proteins among Group X, Y and Z

**Supplementary Table 5**. Overview of B12 biosynthesis protein expression among Group X, Y and Z

The table lists UniProt protein ID, X value representing Log2 Protein abundance ratio (Rhamnose *pdu* BMC induced/non-induced in MWB), Y value representing -Log10 p-value (Rhamnose *pdu* BMC induced/non-induced in MWB), NCBI protein Annotation and NCBI Gene ID.

## References

1. Prevention, C.f.D.C.a., Quantitative assessment of the relative risk to public health from foodborne Listeria monocytogenes among selected categories of ready-to-eat foods. 2003, US Department of Agriculture–Food Safety and Inspection Service Washington, DC.

2. Freitag, N.E., G.C. Port, and M.D. Miner, Listeria monocytogenes—from saprophyte to intracellular pathogen. Nature Reviews Microbiology, 2009. 7(9): p. 623–628.

3. Radoshevich, L. and P. Cossart, Listeria monocytogenes: towards a complete picture of its physiology and pathogenesis. Nature Reviews Microbiology, 2018. 16(1): p. 32.

4. Gandhi, M. and M.L. Chikindas, Listeria: a foodborne pathogen that knows how to survive. International journal of food microbiology, 2007. 113(1): p. 1–15.

5. Portnoy, D.A., V. Auerbuch, and I.J. Glomski, The cell biology of Listeria monocytogenes infection: the intersection of bacterial pathogenesis and cell-mediated immunity. The Journal of cell biology, 2002. 158(3): p. 409–414.

6. Mengaud, J., et al., E-cadherin is the receptor for internalin, a surface protein required for entry of L. monocytogenes into epithelial cells. Cell, 1996. 84(6): p. 923–932.

7. Tattoli, I., et al., Listeria phospholipases subvert host autophagic defenses by stalling pre-autophagosomal structures. The EMBO journal, 2013. 32(23): p. 3066–3078.

8. Kim, H. and A.K. Bhunia, Secreted Listeria adhesion protein (Lap) influences Lap-mediated Listeria monocytogenes paracellular translocation through epithelial barrier. Gut pathogens, 2013. 5(1): p. 1–11.

9. Burkholder, K.M. and A.K. Bhunia, Listeria monocytogenes uses Listeria adhesion protein (LAP) to promote bacterial transepithelial translocation and induces expression of LAP receptor Hsp60. Infection and immunity, 2010. 78(12): p. 5062–5073.

10. Desai, A.N., et al., Changing epidemiology of Listeria monocytogenes outbreaks, sporadic cases, and recalls globally: A review of ProMED reports from 1996 to 2018. International Journal of Infectious Diseases, 2019. 84: p. 48–53.

11. Tasara, T. and R. Stephan, Cold stress tolerance of Listeria monocytogenes: a review of molecular adaptive mechanisms and food safety implications. Journal of food protection, 2006. 69(6): p. 1473–1484.

12. Zeng, Z., et al., Bacterial Microcompartments Coupled with Extracellular Electron Transfer Drive the Anaerobic Utilization of Ethanolamine in Listeria monocytogenes. mSystems, 2021. 6(2).

13. Zeng, Z., et al., Anaerobic Growth of Listeria monocytogenes on Rhamnose Is Stimulated by Vitamin B(12) and Bacterial Microcompartment-Dependent 1,2-Propanediol Utilization. mSphere, 2021: p. e0043421.

14. Kerfeld, C.A., et al., Bacterial microcompartments. Nature Reviews Microbiology, 2018.

15. Yeates, T.O., C.S. Crowley, and S. Tanaka, Bacterial microcompartment organelles: protein shell structure and evolution. Annual review of biophysics, 2010. 39: p. 185–205.

16. Liu, L.N., Bacterial metabolosomes: new insights into their structure and bioengineering. Microbial Biotechnology, 2021. 14(1): p. 88–93.

17. Jakobson, C.M. and D. Tullman-Ercek, Dumpster diving in the gut: bacterial microcompartments as part of a host-associated lifestyle. PLoS pathogens, 2016. 12(5): p. e1005558.

18. Petit, E., et al., Involvement of a bacterial microcompartment in the metabolism of fucose and rhamnose by Clostridium phytofermentans. PLoS One, 2013. 8(1): p. e54337.

19. Anast, J.M., T.A. Bobik, and S. Schmitz-Esser, The Cobalamin-dependent gene cluster of Listeria monocytogenes: implications for virulence, stress response, and food safety. Frontiers in microbiology, 2020. 11.

20. Zeng, Z., et al., Bacterial microcompartment-dependent 1, 2–propanediol utilization stimulates anaerobic growth of Listeria monocytogenes EGDe. Frontiers in Microbiology, 2019. 10: p. 2660.

21. Prentice, M.B., Bacterial microcompartments and their role in pathogenicity. Current Opinion in Microbiology, 2021. 63: p. 19–28.

22. Schneebeli, R. and T. Egli, A defined, glucose-limited mineral medium for the cultivation of Listeria. Applied and environmental microbiology, 2013: p. AEM. 03538-12.

23. Vizcaino, J.A., et al., 2016 update of the PRIDE database and its related tools (vol 44, pg D447, 2016). Nucleic Acids Research, 2016. 44(22): p. 11033–11033.

24. Oliveira, M., et al., Pathogenic potential of Salmonella Typhimurium DT104 following sequential passage through soil, packaged fresh-cut lettuce and a model gastrointestinal tract. International journal of food microbiology, 2011. 148(3): p. 149–155.

25. Koomen, J., et al., Gene profiling-based phenotyping for identification of cellular parameters that contribute to fitness, stress-tolerance and virulence of Listeria monocytogenes variants. International journal of food microbiology, 2018. 283: p. 14–21.

26. Hulsen, T., J. de Vlieg, and W. Alkema, BioVenn–a web application for the comparison and visualization of biological lists using area-proportional Venn diagrams. BMC genomics, 2008. 9(1): p. 1–6.

27. Szklarczyk, D., et al., STRING v11: protein–protein association networks with increased coverage, supporting functional discovery in genome-wide experimental datasets. Nucleic acids research, 2019. 47(D1): p. D607–D613.

28. Mellin, J., et al., A riboswitch-regulated antisense RNA in Listeria monocytogenes. Proceedings of the National Academy of Sciences, 2013. 110(32): p. 13132–13137.

29. Salazar, J.K., et al., PrfA-like transcription factor gene lmo0753 contributes to L-rhamnose utilization in Listeria monocytogenes strains associated with human food-borne infections. Applied and environmental microbiology, 2013. 79(18): p. 5584–5592.

30. Reniere, M.L., A.T. Whiteley, and D.A. Portnoy, An in vivo selection identifies Listeria monocytogenes genes required to sense the intracellular environment and activate virulence factor expression. PLoS pathogens, 2016. 12(7): p. e1005741.

31. Raimann, E., et al., The alternative sigma factor σL of L. monocytogenes promotes growth under diverse environmental stresses. Foodborne pathogens and disease, 2009. 6(5): p. 583–591.

32. Liu, Y., et al., Home alone: elimination of all but one alternative sigma factor in Listeria monocytogenes allows prediction of new roles for σB. Frontiers in microbiology, 2017. 8: p. 1910.

33. Lungu, B., S. Ricke, and M. Johnson, Growth, survival, proliferation and pathogenesis of Listeria monocytogenes under low oxygen or anaerobic conditions: a review. Anaerobe, 2009. 15(1-2): p. 7–17.

34. Scortti, M., et al., The PrfA virulence regulon. Microbes and Infection, 2007. 9(10): p. 1196–1207.

35. Stewart, A.M., et al., Advances in the World of Bacterial Microcompartments. Trends in Biochemical Sciences, 2021.

36. Kennedy, N.W., et al., Bacterial microcompartments: tiny organelles with big potential. Current Opinion in Microbiology, 2021. 63: p. 36–42.

37. Liu, L.-N., et al., Protein stoichiometry, structural plasticity and regulation of bacterial microcompartments. Current Opinion in Microbiology, 2021. 63: p. 133–141.

38. Lebreton, A. and P. Cossart, RNA-and protein-mediated control of Listeria monocytogenes virulence gene expression. RNA biology, 2017. 14(5): p. 460–470.

39. de las Heras, A., et al., Regulation of Listeria virulence: PrfA master and commander. Current opinion in microbiology, 2011. 14(2): p. 118–127.

40. Nadon, C.A., et al., Sigma B contributes to PrfA-mediated virulence in Listeria monocytogenes. Infection and immunity, 2002. 70(7): p. 3948–3952.

41. Pentecost, M., et al., Listeria monocytogenes invades the epithelial junctions at sites of cell extrusion. PLoS pathogens, 2006. 2(1): p. e3.

42. Drolia, R., et al., Listeria adhesion protein induces intestinal epithelial barrier dysfunction for bacterial translocation. Cell host & microbe, 2018. 23(4): p. 470-484. e7.

43. Chelakkot, C., J. Ghim, and S.H. Ryu, Mechanisms regulating intestinal barrier integrity and its pathological implications. Experimental & molecular medicine, 2018. 50(8): p. 1–9.

44. Schardt, J., et al., Comparison between Listeria sensu stricto and Listeria sensu lato strains identifies novel determinants involved in infection. Scientific reports, 2017. 7(1): p. 17821.

